# Age-dependent Effect Imposed by Rotenone Exposure

**DOI:** 10.1101/2021.11.01.466812

**Authors:** Madeleine C. Moseley, Ashley Rawls, Valerie M. Sponsel, Mitchel S. Berger, Adam R. Abate, Chin-Hsing Annie Lin

**Affiliations:** Department of Biology, University of Texas at San Antonio, San Antonio, Texas; Department of Molecular Cellular Integrative Neuroscience, Colorado State University, Fort Collins, Colorado; Department of Integrative Biology, University of Texas at San Antonio, San Antonio, Texas; Department of Neurological Surgery, University of California at San Francisco, San Francisco, California; Department of Bioengineering and Therapeutic Sciences, University of California at San Francisco, San Francisco, California; Neuroscience Institute, University of Texas at San Antonio, San Antonio, Texas

## Abstract

**Background:** The increasing awareness that environmental exposure may lead to sporadic neurological disorders has implicated rotenone to the etiology of some neurodegenerative diseases. However, the risk associated with rotenone toxicity remains controversial as a limited amount of research has studied its effects on brain health.

**Objectives:** This work assessed the risk of rotenone exposure to mice of different ages, gender, and duration by examining *in vivo* effects on brains.

**Methods:** Using a mouse model, the impact of rotenone exposure was determined by analyzing the cellular phenotype in the murine brain.

**Results:** Our results highlight the neurological susceptibility to long-term rotenone exposure in younger ages. For such, younger mice exhibit seizures and convulsions, resulting in shorter lifespan. At the cellular level, rotenone exposure specifically alters the migrating neuroblast populations in the dentate gyrus and causes disorganized pyramidal neurons in the CA3 within the hippocampus. Our findings, albeit the absence of transgenerational inheritance, demonstrated age-related outcomes from rotenone exposure.

**Discussion:** We demonstrated that rotenone exposure specifically influences the population of neuroblasts and pyramidal neurons residing in the hippocampus, a brain region important for learning/memory and associated with convulsive seizure. Our understanding of how exactly rotenone affected region-specific neuronal cells and the molecular mechanism behind exposure risk is still limited. From the perspective of public health, our *in vivo* study highlights age-related susceptibility to rotenone toxicity. Future investigations in environmental epidemiology should determine whether age and duration of exposure to rotenone in human subjects pertains to the development of seizures or other neurological abnormalities over time.

## INTRODUCTION

Environmental exposures have been implicated in the etiology of neurological diseases (Casida, 2009; Caudle et al., 2012; Hatcher et al., 2008; Litvan et al., 2016; Malek et al., 2015; Nandipati and Litvan, 2016; Pearson et al., 2016; Richardson et al., 2014; Richardson et al., 2015; Sanchez-Santed et al., 2016) and developmental defects (Richardson et al., 2015; Shelton et al., 2014). Rotenone, a compound extracted from roots of the *Leguminosae* plant family, is used as a pesticide for agricultural purposes (Dalu et al., 2015). A characteristic of rotenone is that it is lipophilic, and can easily cross the blood-brain barrier and biological membranes. Studies have determined that rotenone causes cell death through the inhibition of mitochondrial electron transport. Rotenone reduces the transfer of electrons from complex I to ubiquinone and limits the activity of mitochondrial NADPH dehydrogenase (Cabeza-Arvelaiz and Schiestl, 2012; Heinz et al., 2017; Tanner et al., 2011). Consequently, oxidative phosphorylation is decreased, resulting in the depletion of ATP from mitochondria. This phenomenon was suspected to be associated with the loss of nigrostriatal dopaminergic neurons and subsequent depletion of dopamine release in the striatum (Cannon et al., 2009). Therefore, rotenone exposure has been linked to sporadic cases of Parkinson’s disease (PD) (Betarbet et al., 2000; Tanner et al., 2011). Despite laboratory research on rotenone in the context of PD, the limitation of this rotenone model is that it does not precisely reproduce the features of PD (Betarbet et al., 2000; Cannon et al., 2009; Pan-Montojo et al., 2010). Besides the rodent model, previous study using a rotenone-treated neuroblastoma cell line has shown that genes involved in the cell cycle were affected (Cabeza-Arvelaiz and Schiestl, 2012). Nonetheless, the effect of rotenone exposure on neurogenesis remains undetermined.

Neurogenesis is the process by which neurons are generated, and the maintenance of neural stem progenitor populations throughout life (Jessberger and Gage, 2014; Kempermann, 2014). After completing brain development, neurogenesis predominately occurs in two adult neurogenic niches: the subventricular zone (SVZ) adjacent to the lateral ventricle (LV) and the subgranular zone (SGZ) of the dentate gyrus (DG) within the hippocampus (Andersen et al., 2014; Ernst et al., 2014; Gallo and Deneen, 2014; Gotz and Huttner, 2005; Ihrie and Alvarez-Buylla, 2011; Khalaf-Nazzal and Francis, 2013; Martynoga et al., 2012; Ming and Song, 2011; Paridaen and Huttner, 2014; Rowitch and Kriegstein, 2010). During neurogenesis within the SVZ, radial glial-like (RGL) cells expressing glial fibrillary acid protein (GFAP) become Nestin^+^ intermediate progenitor cells, which subsequently become doublecortin^+^ (DCX^+^) microtubule-associated migrating neuroblasts. DCX^+^ neuroblasts eventually migrate out of the SVZ via the rostral migratory stream and differentiate into neurons in the olfactory bulb (Bond et al., 2015; Ihrie and Alvarez-Buylla, 2011; Kempermann et al., 2015; Ming and Song, 2011). Likewise, during neurogenesis within the SGZ of DG, GFAP^+^ RGL cells give rise to Nestin^+^ intermediate progenitors and then DCX^+^ neuroblasts, which differentiate into mature neurons in the DG (Andersen et al., 2014; Bond et al., 2015; Ernst et al., 2014; Kempermann et al., 2015; Ming and Song, 2011). Mature neurons receive excitatory signals and project the signals along the mossy fiber tract to cornu ammonis 3 (CA3) during their maturation phase. The signal from CA3 projects to cornu ammonis 1 (CA1) that then allows for reverse projection back to the neocortex for the retrieval of information. Thus, CA3, CA1, and the DG are a part of the trisynaptic circuit of adult neurogenesis within the SGZ/DG (Andersen et al., 2014; Ernst et al., 2014; Hunsaker et al., 2006; Kempermann et al., 2015; Ming and Song, 2011). Importantly, the intimate relationship between the cell fate of undifferentiated cells in the SGZ/DG and the lamination of pyramidal neurons in the CA3 is implicated in neuropathological conditions, such as convulsion, epilepsy, and seizure (Nosten-Bertrand et al., 2008). Previous studies to understand these neurological conditions associated with the hippocampus were mostly concerned with genetic factors and signaling pathways. The outcome of environmental exposure on these types of neuropathological conditions remains unexplored.

Our study on rotenone exposure in a murine model showed that DCX^+^ neuroblasts were decreased specifically in the DG while no obvious changes in either dividing or differentiated cells were detected, suggesting neurogenesis was not compromised upon rotenone exposure. However, the defect in DCX^+^ migrating neuroblasts contributed to disorganized pyramidal layer of the CA3 within the hippocampus. Intriguingly, we found that younger mice were more vulnerable to rotenone exposure and showed profound phenotypes including hyperactivity and seizure-associated early death. These behavioral and cellular phenotypes were not present in offspring (F1), suggesting the negative effects of parental rotenone exposures are reset in the subsequent generation. In the perspective of public health implication, our *in vivo* study highlights age-related susceptibility to rotenone toxicity.

## METHODS

### Animal Models

The strains of mice used for this study were C57BL6J and C57BL6J/129SV background including both male and female. The analysis includes different ages of mice from 1 month to 10 months old to recapitulate the age of exposure in human beings. All animal experiments were approved by the Institutional Animal Care and Use Committee of the University of Texas at San Antonio (UTSA).

### Intraperitoneal Injections

The rotenone stock solution (15 mg/ml) was prepared by dissolving 15 mg Rotenone (Sigma Aldrich, Lot#MKBz2534V) in 1 ml DMSO (dimethyl sulfoxide). The rotenone working solution was freshly prepared with 20 microliters (μl) of stock solution and 980 μl of Miglyol 812 N (IOI Oleochemical, Lot#170709) or Sunflower Oil. Based on body weight, a total of 0.5-1 ml of rotenone working solution was injected per each mouse (3mg/kg body weight). Mice were subjected to daily rotenone exposure via intraperitoneal injection for 20 days and 40 days. Untreated control mice were littermate without injection while vehicle control mice were injected with Miglyol 812N or Sunflower Oil.

### Transcardial Perfusion and Fix of Mice

The mice were anesthetized with Avertin (450 mg/kg) before perfusion/fixation transcardially with 30 ml of 1x PBS and 4% paraformaldehyde. Following post-fixation for up to an hour in 4% paraformaldehyde, brains were cryopreserved with 30% sucrose/PBS overnight and frozen in OCT compound. Cryo-protected brains were OCT embedded, and 14-μm frozen sections were cut for Nissl staining (every 5 sections). Slides undergoing Nissl Stain were incubated in Thionine for 10 seconds then rinsed in water 3 times. After water rinse, slides were subsequently dipped in 70%, 95%, and 100% ethanol 5 times. After ethanol, slides were dipped in Xylene before mounting the coverslip. Before applying the coverslip, Shandon Consul-Mount Xylene base was added to each section.

### Histochemistry and Immunostaining

Frozen brain sections on super frost microscope slides were rinsed with 1x PBS until OCT compound dissolved from the slides. Each section was then incubated for 0.5 hour with 10% goat serum in 1x PBST (1x PBS and 0.3% TritonX-100) for blocking. After blocking for 30 minutes, primary antibodies were incubated with sections at 4 °C overnight. Antibodies specific for cell type specific markers: GFAP (Millipore, Clone GA5, Cat#mab3402, Lot#2580632, 1:750 and Invitrogen, Lot#TF268267, 1:500), NeuN (Millipore, clone A60, Cat#MAB377, Lot#NG1876252, 1:200), TH (Millipore, Lot#2950733, 1:200), DCX (Santa Cruz, Cat#SC-8066, Lot#10314, 1:500), Nestin (Millipore, Cat#MAB353, 1:200), Ki67 (Cell Signaling, #9449S, 1:200), Tuj1 (Millipore #MAB1637, 1:200), P-H3 (Millipore #06-570; Cell Signaling #9307S, 1:200), NPY (Bioss #bs0071R, 1:200), and Calbindin (Abcam #ab82812, 1:200). The following day, sections were washed 3 times with PBST before incubation with fluorescent-conjugated secondary antibodies (1:500 in PBST) for 1 hour at 4°C, protected from light. AlexaFlour-488 or -594 conjugated secondary antibodies (anti-mouse: Life Technologies, Cat#A21200; Lot#1776045; anti-rabbit: Life Technologies, Cat#A21207; Lot#1668552) were added to each section that coordinated with primary antibodies. The Thymidine analog incorporation for analysis of dividing cells was done through intraperitoneal injections of EdU (5-ethynl-2’-deoxyuridine) at the concentration of 150 μg/g of body weight per mouse 72 hours prior to perfusion/fixation. EdU-positive cells were detected using Click-iT EdU 488 imaging kit (Invitrogen, C10340) on cryo-protected brain sections following the manufacturer’s instructions. All of the sections were mounted with Vectashield mounting medium with DAPI (Vector Laboratories, Cat# H-1200). Images were acquired on Zeiss 710 confocal microscope with 10x, 20x, and 40x objectives. Images were processed using the Zen software and scale bars were added using Image J https://imagej.nih.gov/ij/). Quantification: Raw confocal Z-stacks images were uploaded to ImageJ, converted to 16-bit grayscale, and then a global automatic threshold was applied to each image. Cell counts of DCX were normalized to DAPI numbers of the entire image to present percentage (%) in the DG. All data were plotted as mean % ±SD (standard deviation), and the significances were determined by two-way ANOVA with the Tukey’s post-hoc test.

## RESULTS

### Age susceptibility to rotenone exposure

Behavioral phenotype is one readout of brain function, which encompasses various factors like age and gender. To determine if age and gender play roles in the susceptibility to rotenone exposure, the relationship of the stages of growth in mice to human age was used (Wang et al., 2020). The timing of growth and developmental stages, including suckling period, juvenile period, puberty, and adulthood, can be used to make the correlation between human years and age of mice in weeks. Time points of exposure in relation to duration in humans were also calculated using the relationship of lifespan between mice and humans, in which a 2 year-old mouse is roughly equivalent to an 80 year-old human (Dutta and Sengupta, 2016). In this regard, the various ages of mice tested were between 1-2 month of age (n=42), 3-4 months (n=22), and 5-10 months (n=26) of age. We utilized daily intraperitoneal injection (IP) into adult mice with a range of doses (0.5 - 3mg/kg body weight), and then focused on the maximum dose of 3mg/kg body weight for 20- and 40-day treatments (Betarbet et al., 2000; Cannon et al., 2009). Different solvents were used in previous rotenone studies (Betarbet et al., 2000; Cannon et al., 2009; Wang et al., 2018) including Miglyol and sunflower oil. Miglyol is composed of C6-C12 medium chain and C14 long-chain fatty acids while sunflower oil contains a higher percentage of long-chain fatty acids (C14-C18). As there is limited information about the effect of lipids like Miglyol and sunflower oil on the brain, we unbiasedly used both solvents for vehicle controls and dissolving rotenone.

Across age groups, mice exposed to rotenone showed behavioral abnormalities including hyperactivity (Carter and Shieh, 2015) during the first week of exposure. Regardless of the length of exposure, there were more cases of early death combined with convulsion/seizure phenotypes in mice who were 1-2 months old at the initial time of exposure (52%) than in mice that were 3-10 months of age (38%). The difference between convulsions and seizures is that convulsions are instances of repeated short seizures that can be recovered from, while seizures are classified as large incidents that result in death. Our finding suggests a greater susceptibility to rotenone in those that are younger. When observing early death cases in 1-2 months old mice, susceptibility predominately occurs within the first two weeks of exposure and the probability of early death decreases after. Taken together, the critical time point of exposure is within 2-weeks when mice begin to exhibit phenotypes consisting of seizures/convulsions. Results of convulsion/seizure and early death suggest that younger ages are more susceptible to rotenone exposure.

### Neurogenic niches: Rotenone affects DCX migrating neuroblasts in the DG and organization of CA3 cells within the hippocampus

Considerable evidence suggests behavioral phenotypes like seizures are linked to dysregulated neurogenesis in the DG of the hippocampus (Jessberger and Parent, 2015; Nosten-Bertrand et al., 2008). As the process of neurogenesis is aided by cell cycle progression, we first assessed the impact of rotenone exposure on the S-phase of the cell cycle by intraperitoneal injection with 5-ethynyl-2’-deoxyuridine (EdU) for 72 hours tracing (Figure 1). When comparing untreated littermate control (Figure 1A) with vehicle controls, sunflower oil and Miglyol do not impact the population of dividing cells within the DG and the SVZ (Figure 1B, D, Supplemental Figure 1). In 1-month-old mice, rotenone exposure affects the distribution of dividing cells undergoing the S-phase of the cell cycle in cases of convulsion, seizure, postural instability, and rigidity (Figure 1C) but this cellular effect was not seen when rotenone did not induce severe seizure (Figure 1E). Among all age groups tested, rotenone does not alter EdU^+^ cells residing in the SVZ (Supplemental Figure 2), suggesting the DG region-specific effect from rotenone exposure. In contrast to 1-month-old mice, rotenone exposure in mice 2-months and older does not influence the EdU^+^ dividing cells in the DG of the hippocampus and the SVZ, regardless of occurrences of behavioral abnormalities (Supplemental Figure 3, data not shown). Next, with the focus on the DG of the hippocampus, immunostaining of Ki67 and phosphor-histone H3 (PH3) was conducted to examine the population of proliferating cells at interphase and those undergoing mitosis, respectively. Rotenone does not affect the population of interphase and mitotic cells in the DG in both 1-month-old and ≥2 months old mice (Figure 2), suggesting that cell cycle progression was not altered upon rotenone exposure.

**Figure 1:**
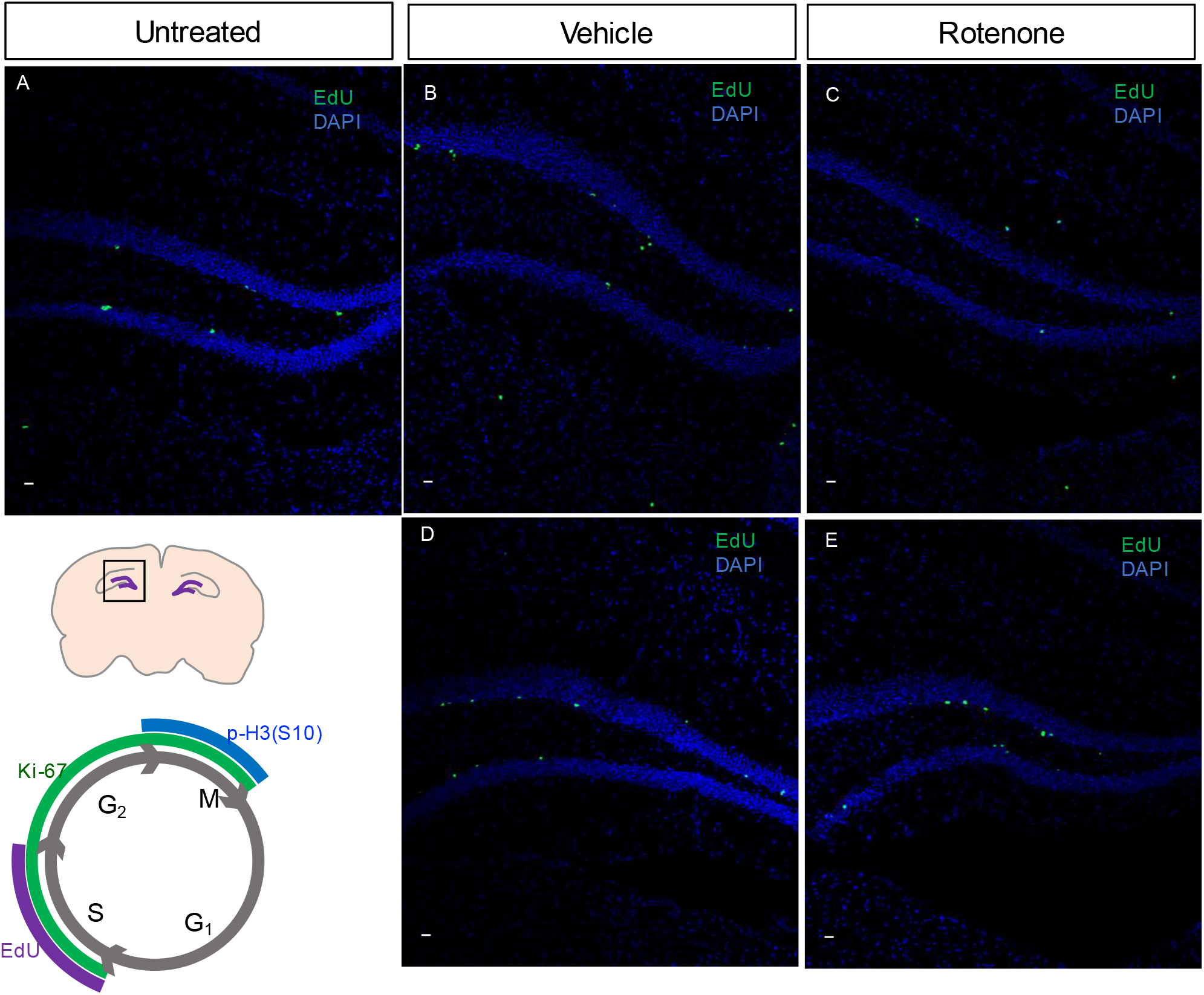
Rotenone exposure-associated behavioral abnormalities showed altered distribution pattern of dividing cells in the DG of the hippocampus in 1-month-old mice. Representative images from 72 hours EdU tracing in 1 month old mice. (A) 1 month old untreated mouse. (B) vehicle control: sunflower oil treated 1 month old mouse. (C) rotenone expose: rotenone in sunflower oil treated 1 month old mouse that experienced convulsions, postural instability, and rigidity. (D) vehicle control: sunflower oil treated 1 month old mouse. (E) Rotenone expose: rotenone in sunflower oil treated 1 month old mouse that did not experience behavioral abnormalities. (B-C) and (D-E) are littermates. Scale bars = 10 µm.

**Figure 2:**
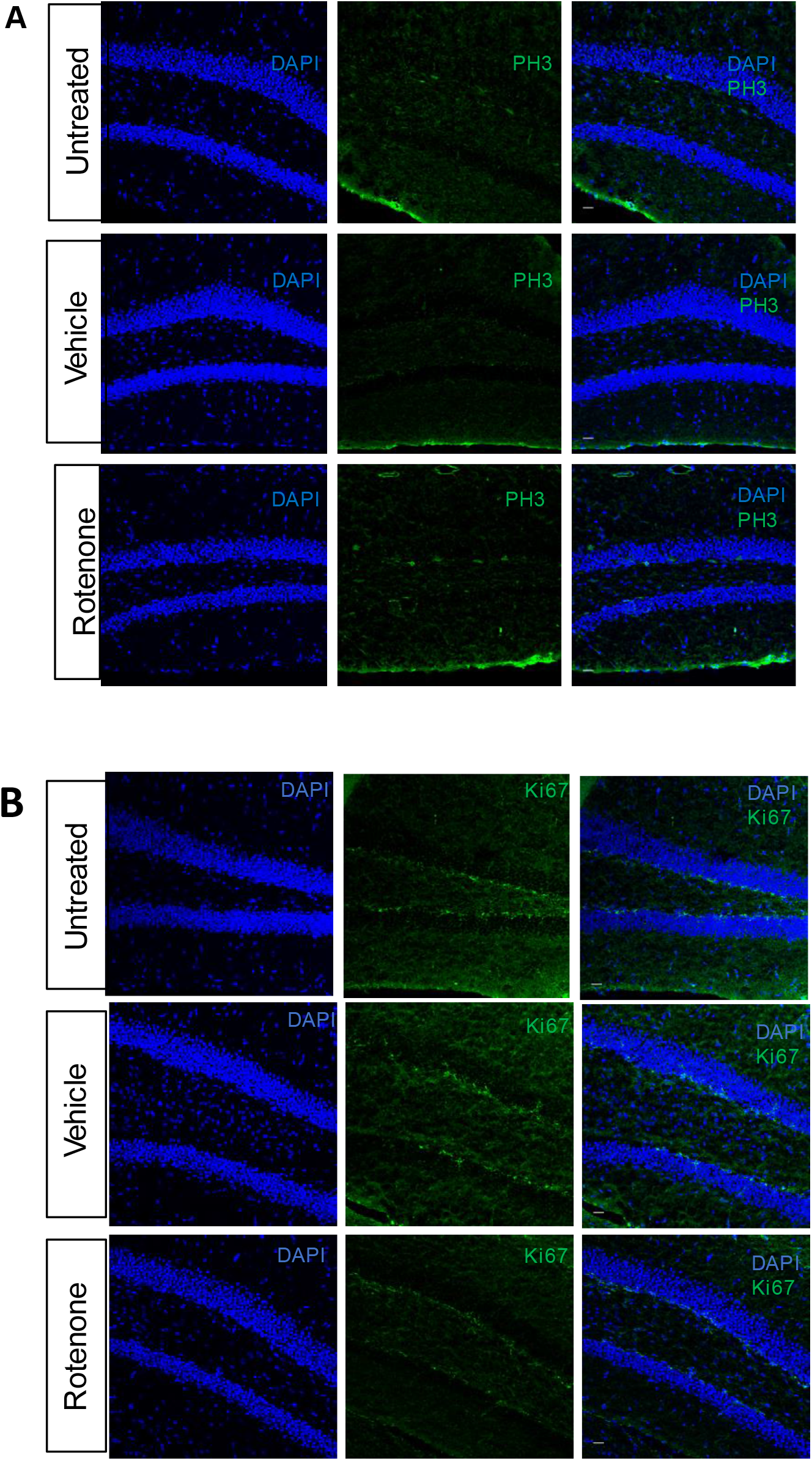
Rotenone exposure does not affect the cell cycle progression. Representative images of (A) PH3^+^ mitotic cells, (B) Ki67^+^ interphase cells in the DG within the hippocampus are not affected after exposure. Scale bars = 20 µm.

Neurogenesis begins with the process when GFAP^+^ RGLs first enter the cell cycle and undergo divisions to give rise to intermediate progenitors and neuroblasts, which shortly exit the cell cycle (G0) becoming early-born postmitotic neurons (Andersen et al., 2014; Ernst et al., 2014; Hunsaker et al., 2006; Kempermann et al., 2015; Ming and Song, 2011). Herein, we utilized cell-type specific markers to assess the impact of rotenone exposure on the population of neural stem progenitor cells. In 1-month and 2-month-old mice, rotenone does not alter GFAP^+^ RGLs (Figure 3) and Nestin^+^ intermediate progenitors (Figure 4), but decreases the population of DCX^+^ neuroblasts in the DG (Figure 3). By contrast, DCX^+^ neuroblasts were unaffected in the SVZ (Supplementary Figure 4), suggesting rotenone has a region-specific effect on neuroblasts in the DG of the hippocampus. However, rotenone does not influence NeuN^+^ mature neurons (Figure 5), pre-mature neurons (Tuj1^+^) (Figure 6A), and Calbindin^+^ interneurons (Figure 6B). Although our results demonstrated that neurogenesis was not affected upon rotenone exposure, Nissl staining showed disorganized pattern of cells within the CA3 (Figure 7A) reminiscent of a publication from a *Dcx* knock-out mouse model, in which a lack of lamination of pyramidal neurons within the CA3 of the hippocampus has been linked to convulsive seizures (Nosten-Bertrand et al., 2008). We utilized the marker neuropeptide Y (NPY) to further demonstrate the defect of pyramidal neurons in the CA3 (Figure 7B).

**Figure 3:**
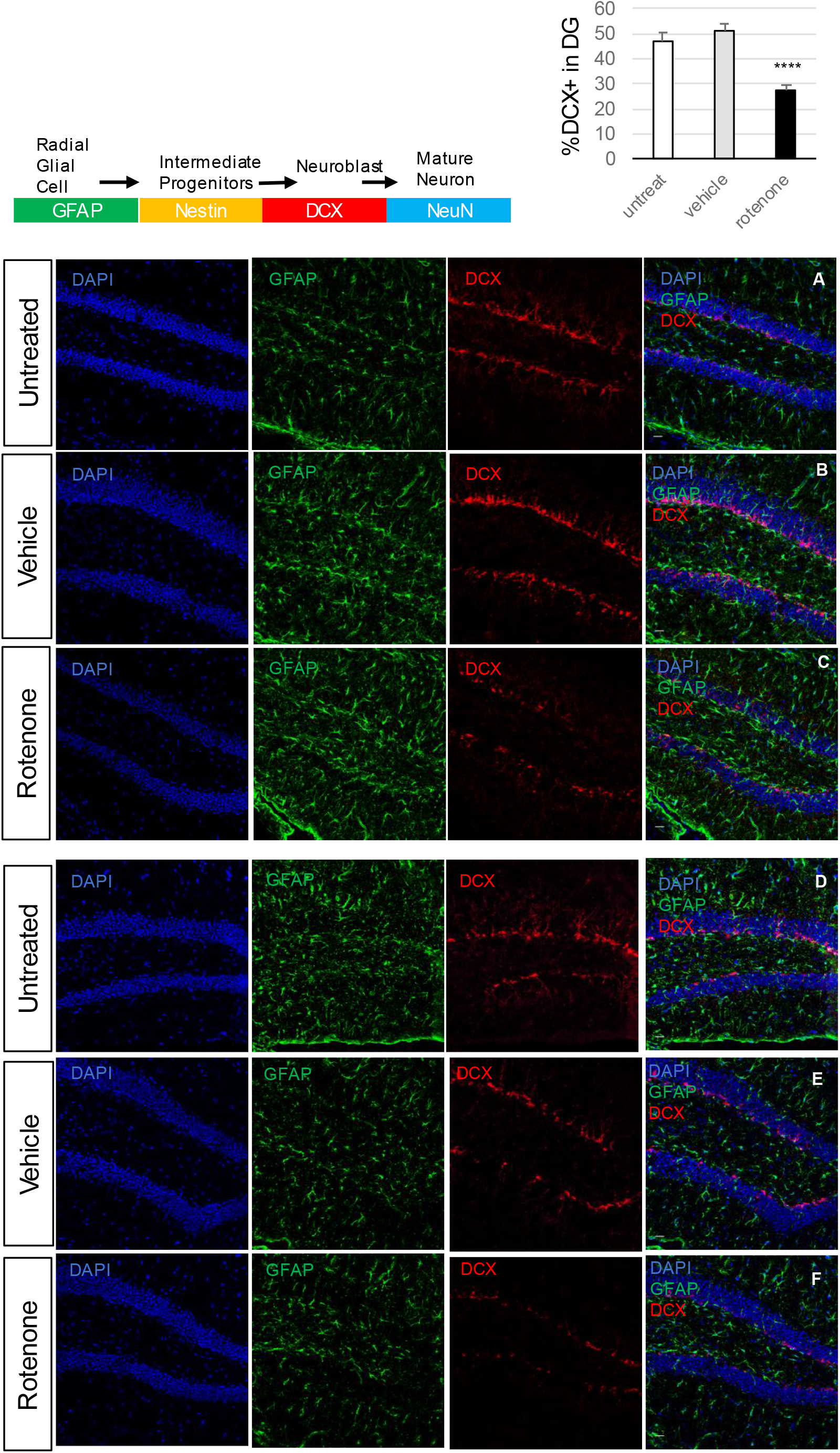
Rotenone exposure affected neuroblasts in the DG within the hippocampus in 1-month and 2-month-old mice. Scheme dictates the stepwise transition from neural stem cell (also known as radial glial-like cells) toward differentiated neurons in the DG of hippocampus. Bar graph summarized average % of DCX^+^ neuroblasts in the DG for untreated, vehicle control, and rotenone exposed mice. Statistical analysis: no significant differences between untreated and vehicle control (p-value = 0.4). A significant difference was found (p-value = 2.39545E-05) when comparing rotenone-treated versus untreated/vehicle while significances were also found in the comparisons of rotenone-treated versus untreated (p-value = 0.0025) and rotenone-treated versus vehicle control (p-value = 0.00017). Representative images show reduction of DCX^+^ neuroblasts and no change in GFAP^+^ RGLs. (A-C) 1-month-old mice. (D-F) 2-month-old mice. (A and D) Untreated mice. (B and E) Sunflower oil treated mice. (C and F) Rotenone in sunflower oil treated mice. Scale bars = 20 µm.

**Figure 4:**
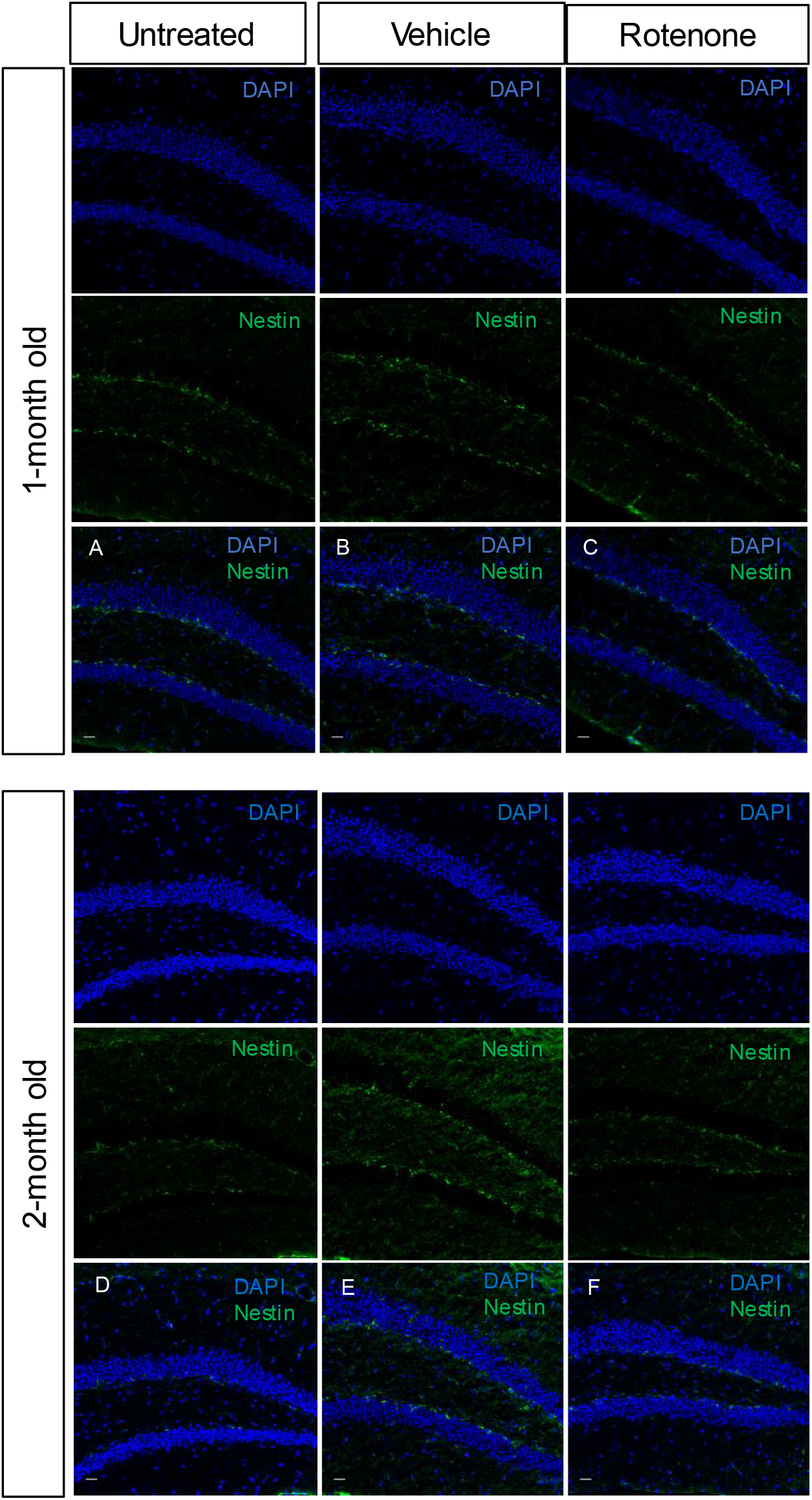
Rotenone exposure does not affect intermediate progenitors in the SGZ of the DG within the hippocampus in 1-month or 2-month-old mice. Representative images of Nestin^+^ intermediate progenitors show no change after exposure. (A-C) 1-month-old mice. (D-F) 2-month-old mice. (A and D) Untreated mice. (B and E) vehicle control: Sunflower oil treated mice. (C and F) Rotenone in sunflower oil treated mice. Scale bars = 20 µm.

**Figure 5:**
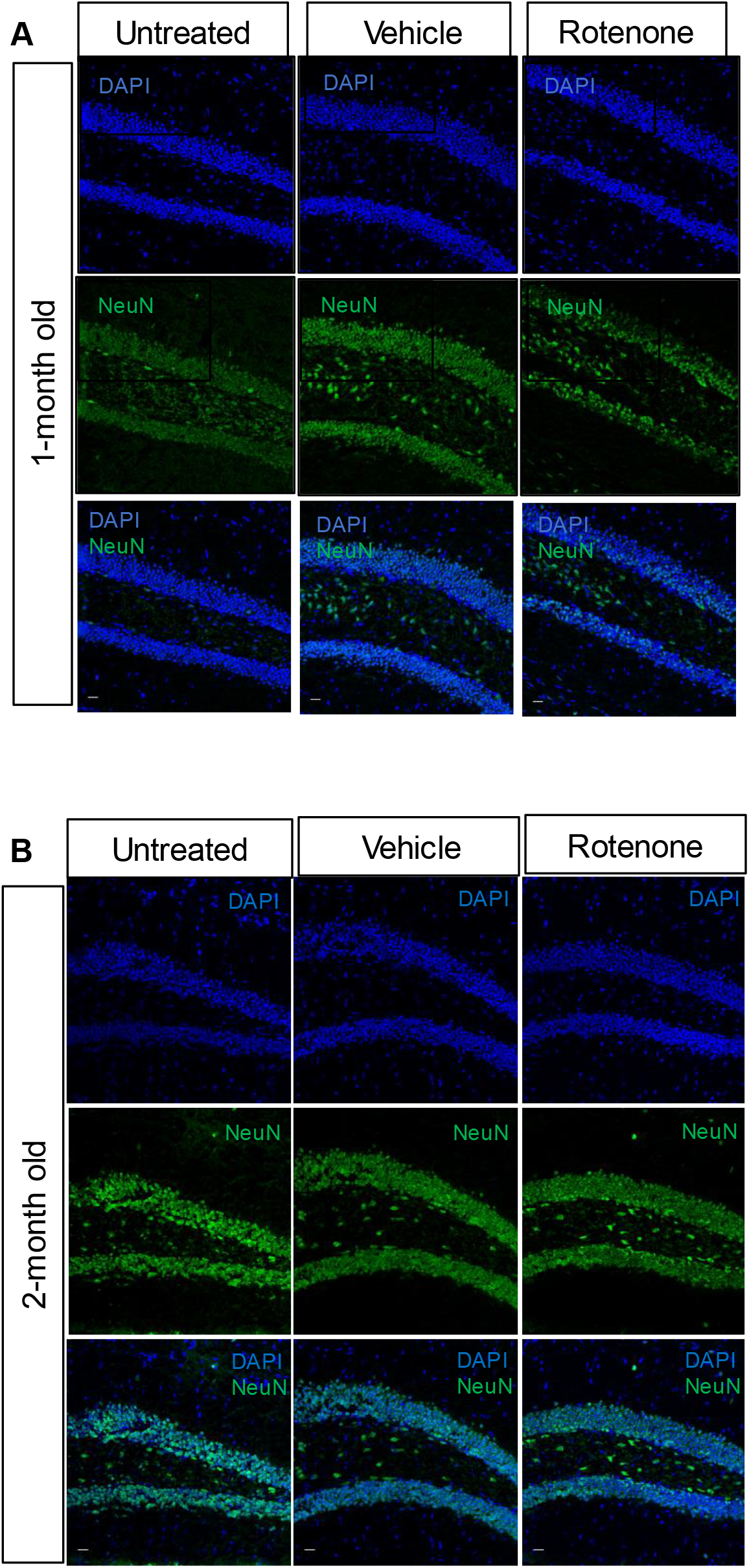
Rotenone exposure does not affect mature neurons in the DG within the hippocampus in 1-month-old and 2-month-old mice. Representative images of NeuN^+^ mature neurons in (A) 1-month-old, and (B) 2-month-old showed rotenone exposure does not affect mature neurons. Scale bars = 20 µm.

**Figure 6:**
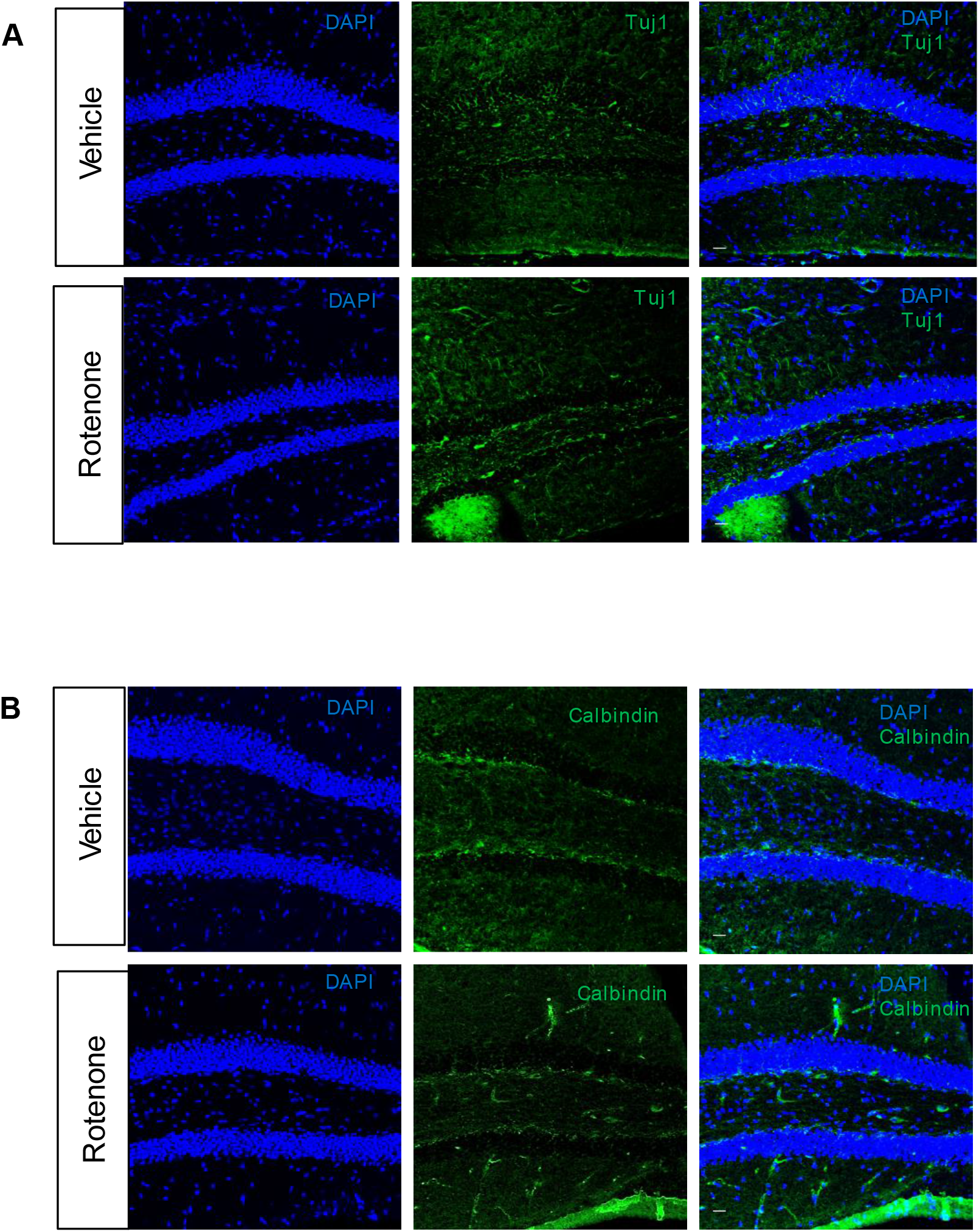
Rotenone exposure does not affect pre-mature neurons and interneurons in the DG within the hippocampus in 1-month-old mice. Representative images of (A) Tuj1^+^ pre-mature neurons, (B) Calbindin^+^ interneurons. Scale bars = 20 µm.

**Figure 7:**
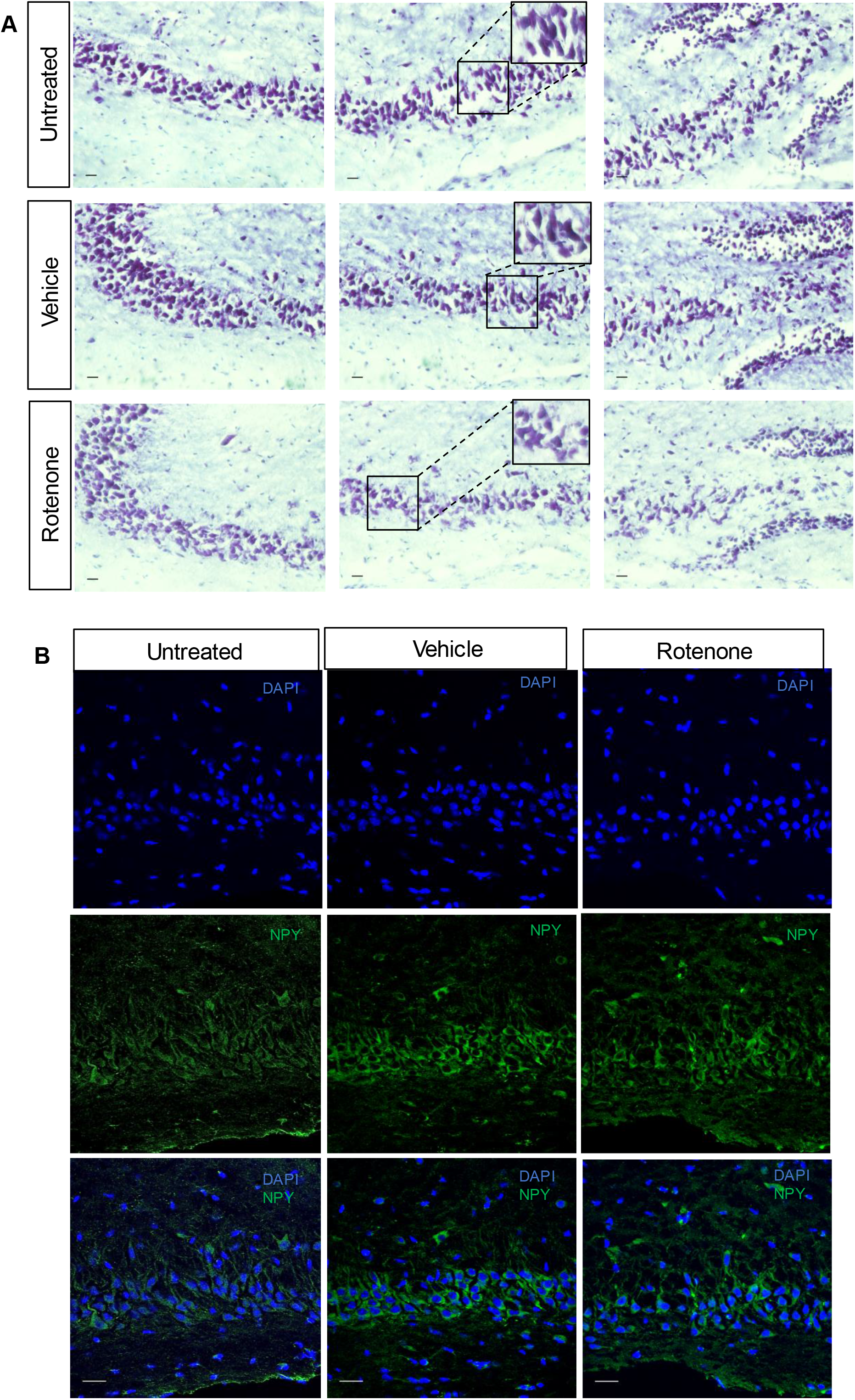
Rotenone exposure affects the organization of pyramidal cells within the CA3 of the hippocampus in 1-month old rotenone-exposed mice. (A) Representative images of Nissl staining showed dispersed patterns especially for cells residing in the CA3b, middle panel with box inset. Left-right panels: CA3a, middle panel CA3b. Scale bars=20 µm. (B) NPY immunostaining in the CA3b region showed the NPY^+^ pyramidal cells were disorganized in rotenone treated mice while untreated and vehicle control littermates had aligned NPY^+^ cell clusters. Scale bars = 40 µm.

The notion that pesticide exposure can sometimes affect multiple generations, known as transgenerational inheritance, prompted our further investigation of F1 animals, with the focus on the most profound phenotype DCX^+^ neuroblasts in the DG of hippocampus. The region-specific effect of DCX^+^ neuroblasts in the DG was observed in the parent line F0 but is not present in the F1 generation (Figure 8), suggesting no transgenerational effect imposed by parental exposure to rotenone.

**Figure 8:**
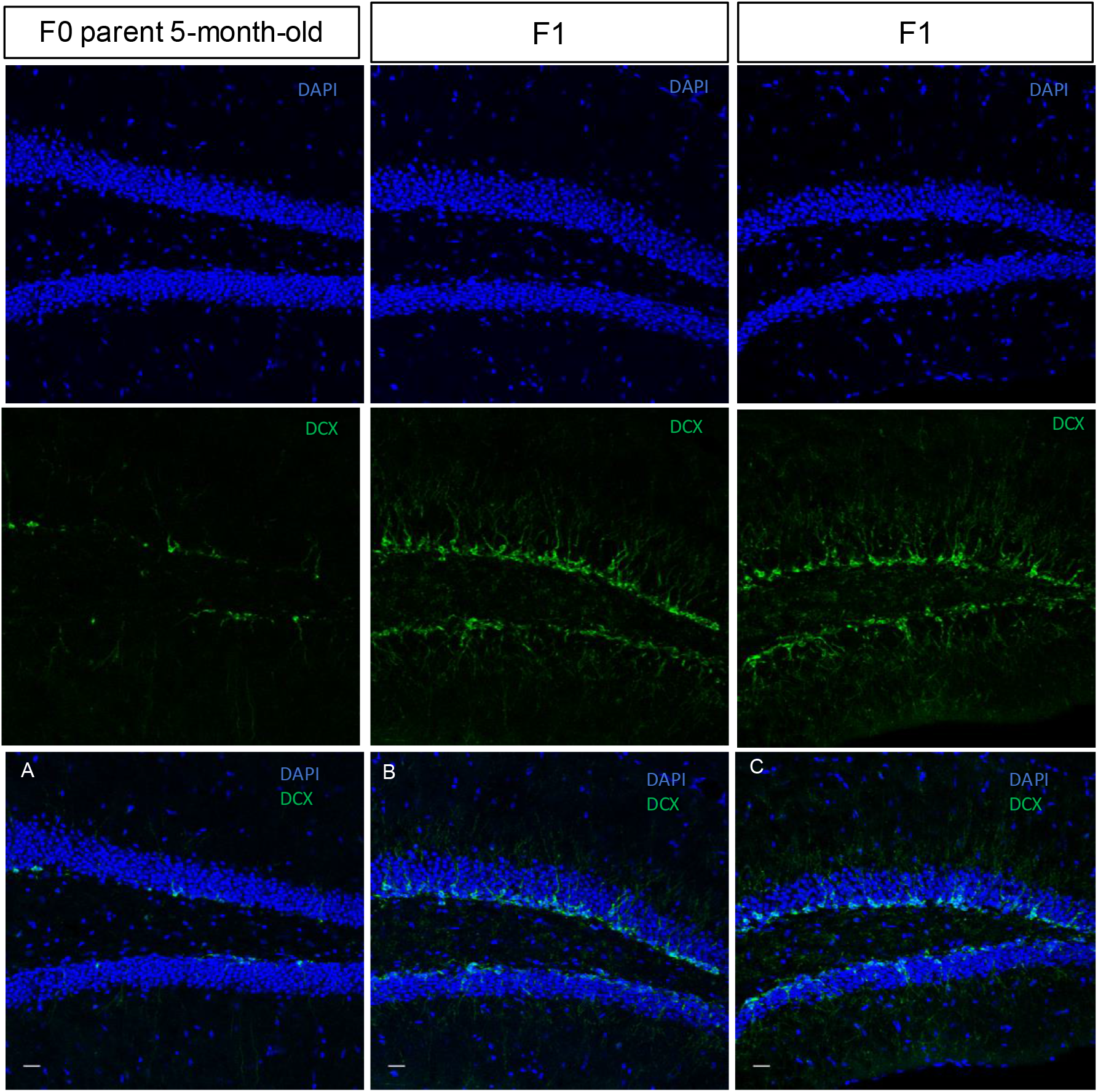
Rotenone’s effect on neuroblasts in the DG within the hippocampus is not transgenerationally inheritable. Representative images of DCX^+^ neuroblasts in the DG of parent (F0) and F1 generation (female and male) showed the cellular phenotype is not inherited to the next generation. (A) F0 parental exposure to rotenone for 20 days. (B-C) 1-month-old F1 generation. Scale bars = 20 µm.

Rotenone exposure has been implicated in sporadic cases of Parkinson’s disease (PD) (Betarbet et al., 2000; Tanner et al., 2011). In PD patients, the dopaminergic neurons and tyrosine hydroxylase (TH) required for dopamine synthesis in the substantia nigra pars compacta (SNc) are degenerated. We examined this cellular phenotype associated with PD and found that TH^+^ neurons in the SNc were not significantly changed after 20- and 40-day rotenone exposure (Supplementary Figure 5).

## DISCUSSION

The use of insecticides and pesticides in industrial agriculture is inevitable, although environmental exposure has been implicated in the etiology of cancer and neurological diseases. As health risks arising from the overuse of insecticides or pesticides has increased public awareness, we assessed rotenone exposure in different age groups, genders, and generations using *in vivo* mouse model. This work presents the first investigation on the impact of rotenone exposure in neurogenic niches. We found that DCX neuroblasts in the DG of hippocampus were affected in exposed mice across age groups, and the pyramidal layer in the CA3 region was disorganized potentially due to decreased or arrested migration of hippocampal neuroblasts. Evidence from *Dcx* knock-out mice showed hyperactivity, and the incidence of epileptic seizures associated with discrete lamination defects and enhanced excitability in the hippocampus (Nosten-Bertrand et al., 2008). Perhaps, rotenone-induced spontaneous seizures underlie the reduction of DCX that consequently affects synaptic transmission involving heterotopic cells in the hippocampus. Given the fact that previous reports had used both miglyol and sunflower oil as solvent to dissolve rotenone, we unbiasedly examined the *in vivo* effect of rotenone in both lipid contents. In vehicle controls, the use of miglyol and sunflower oil did not influence phenotype driven by rotenone exposure. Our understanding of how exactly rotenone affected region-specific neuroblasts is still scarce, and in-depth investigations are needed. Due to the behavioral phenotype dominated by spontaneous convulsive seizures upon rotenone exposure, further study of the molecular consequences of exposure may require an alternative approach, such as the utilization of a brain organoid model.

Our work provides a blueprint for exploratory studies on chemical exposure to determine critical time points and factors (age, gender, length of exposure, and transgenerational inheritance) that offer insight for human exposure risk. In conclusion, we examined the impact of rotenone exposure on brain health to demonstrate that rotenone exposure specifically influences the population of DCX^+^ neuroblasts in the DG, and the organization of pyramidal neurons in the CA3 region within the hippocampus. Future investigations from environmental epidemiology should determine whether age and duration of exposure are factors influencing whether rotenone-exposed human subjects develop seizure or other neurological abnormalities over time.

## ACKNOWLEDGMENT

We thank Angela Huang, David Nweke, and Honor’s students in Lin’s lab for technical assistance. We also thank Dr. Wei-Ming Chien for imaging quantification and statistical analysis. Authors are grateful to Drs. Chris Rhodes, Sandra Cardona, Isabel Muzzio, and Todd Troyer for insightful discussion. Confocal imaging was done in the Cell Analysis Core at UTSA. This project was supported by NIH grant NIH/NIGMS SC3GM112543 and TRAC award to CAL

## FIGURE CAPTIONS

**Supplementary Figure 1:**
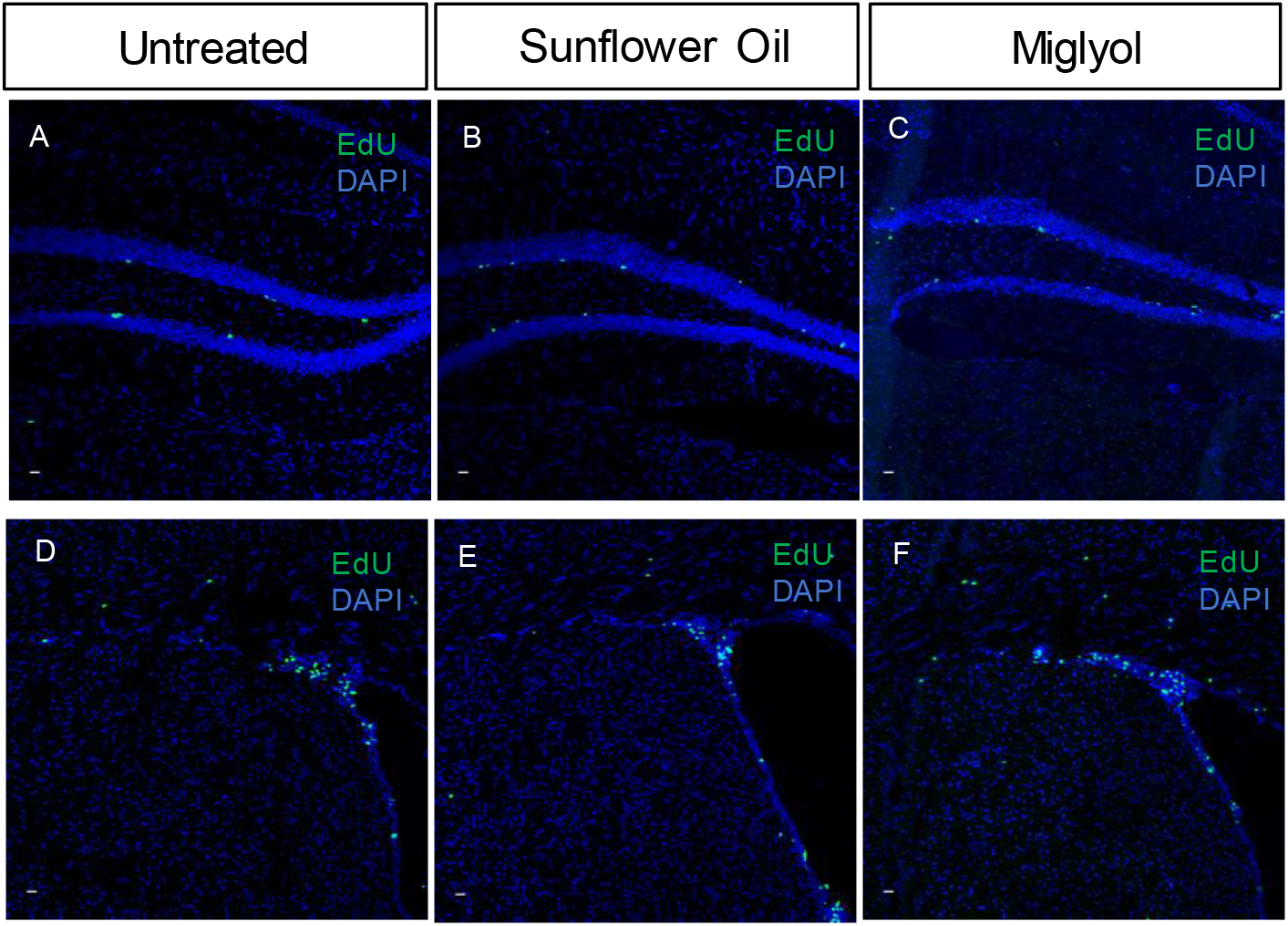
Solvents, sunflower oil and Miglyol, do not affect dividing cells in neurogenic niches. Representative images from 72 hours EdU tracing in 1 month old mice. (A-C) DG within the hippocampus. (D-F) SVZ. (A and D) 1-month-old untreated mouse. (B and E) Vehicle control: Sunflower oil treated 1-month-old mouse. (C-F) vehicle control: Miglyol treated 1-month-old mouse. Scale bars = 10 µm.

**Supplementary Figure 2:**
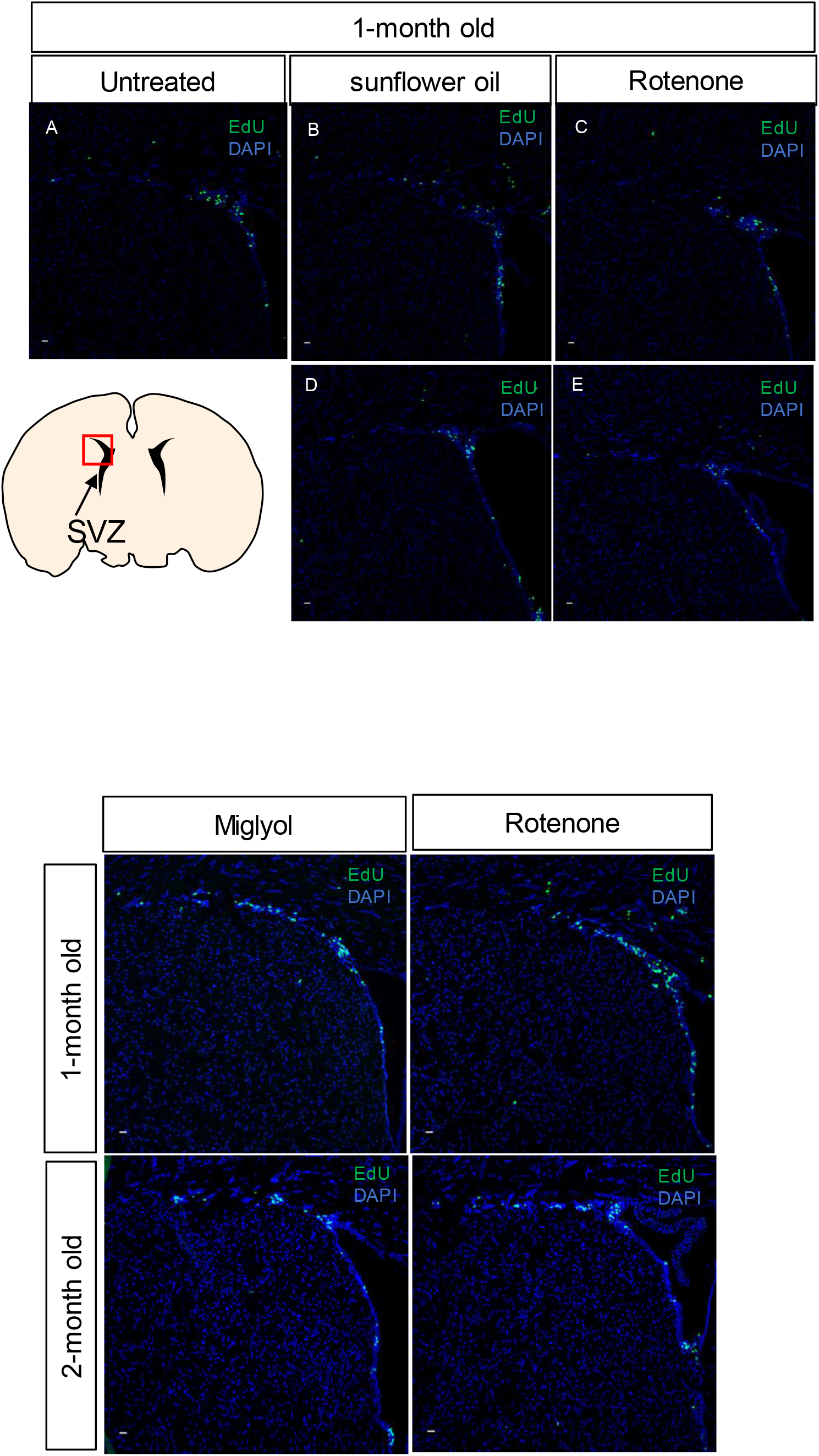
Rotenone exposure did not affect dividing cells in the SVZ. Top Panel: Representative images from 72 hours EdU tracing in 1 month old mice. (A) 1 month old untreated mouse. (B) vehicle control: sunflower oil treated 1 month old mouse. (C) Rotenone in sunflower oil treated 1 month old mouse that experienced convulsions, postural instability, and rigidity. (D) vehicle control: sunflower oil treated 1 month old mouse. (E) Rotenone in sunflower oil treated 1 month old mouse that did not experience behavioral abnormalities. (B-C) and (D-E) are littermates. Scale bars = 10 µm. Bottom panel: Rotenone in Miglyol does not affect dividing cells in neurogenic niches in 1-month or 2-month-old mice. Representative images from 72 hours EdU tracing in 1-month-old and 2-month-old mice. Scale bars = 10 µm.

**Supplementary Figure 3:**
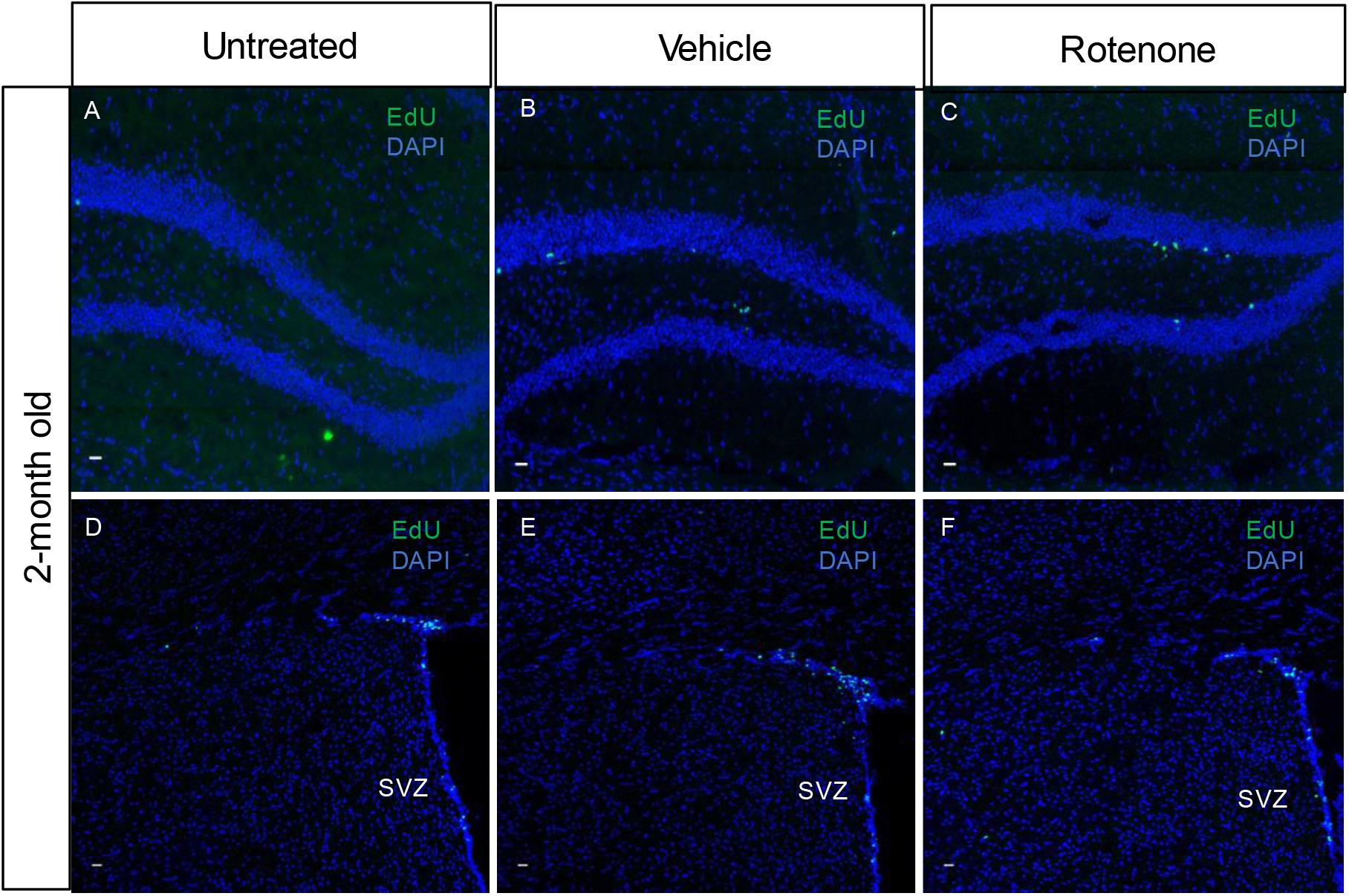
Rotenone exposure does not affect the population of dividing cells in the DG within the hippocampus or the SVZ in 2-month-old mice. Representative images from 72 hours EdU tracing in 2-month-old mice (A-C) DG within the hippocampus. (D-F) SVZ. (A and D) 2-month-old untreated mouse. (B and E) Sunflower oil treated 2-month-old mouse. (C-F) Miglyol treated 2-month-old mouse. Scale bars = 10 µm.

**Supplementary Figure 4:**
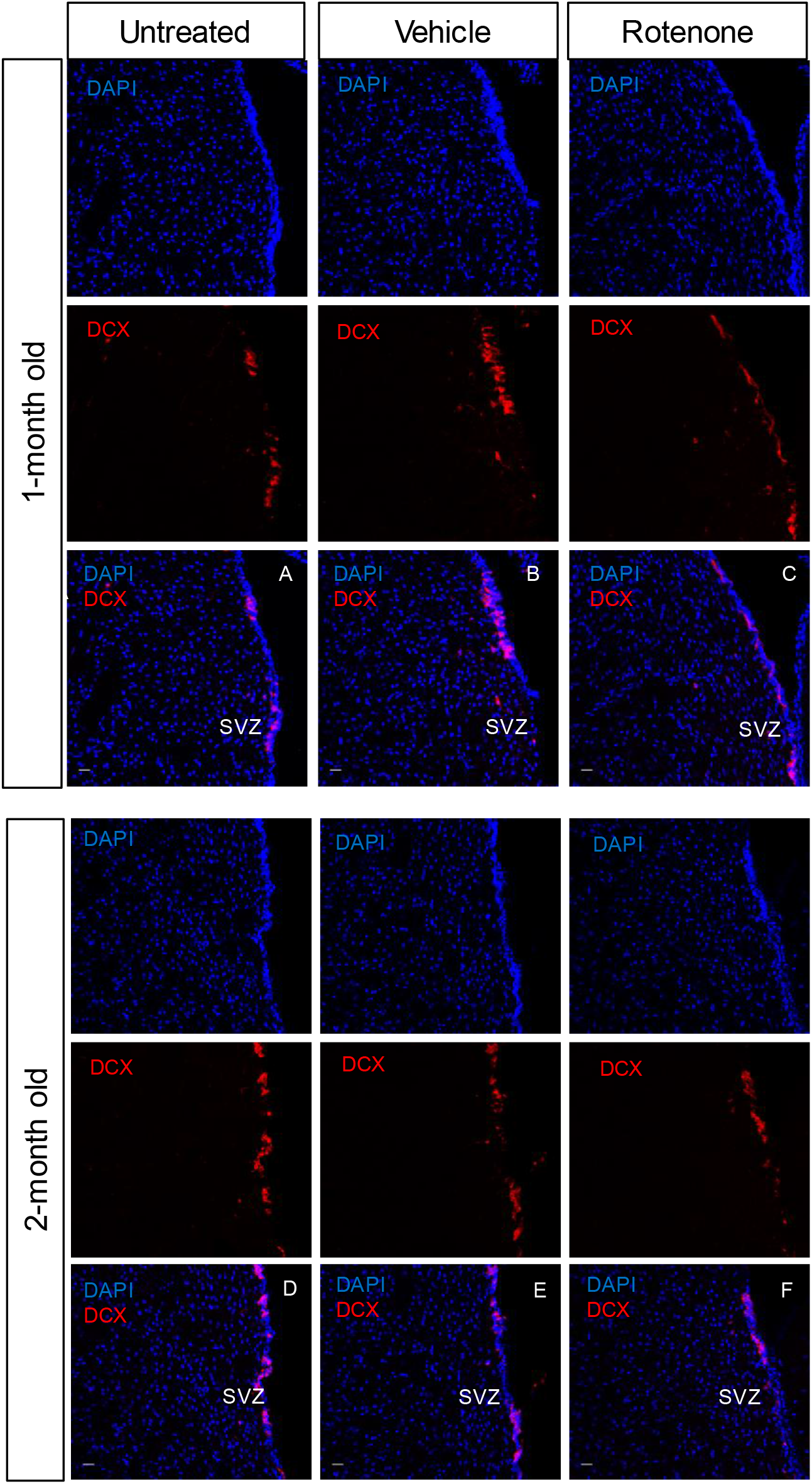
Rotenone exposure does not affect migrating neuroblasts in the SVZ in 1-month or 2-month-old. Representative images of DCX^+^ neuroblasts show no effect by exposure. (A-C) 1-month-old mice. (D-F) 2-month-old mice. (A and D) Untreated mice. (B and E) vehicle control: Sunflower oil treated mice. (C and F) Rotenone in sunflower oil treated mice. Scale bars = 20 µm.

**Supplementary Figure 5:**
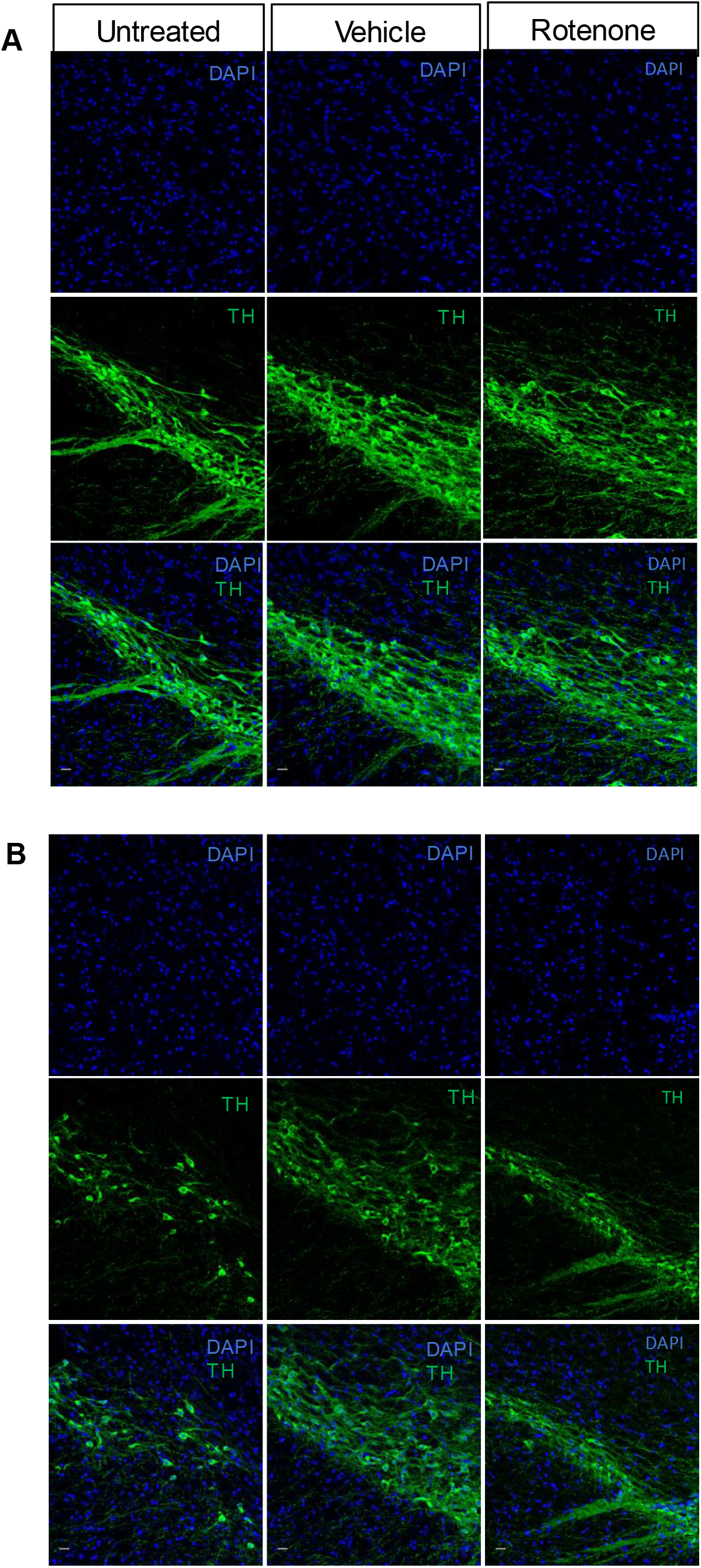
Rotenone exposure does not affect TH^+^ neurons within the SNc in 1-month and 2-month-old mice. Representative images of TH^+^ neurons showed no change by rotenone exposure. (A) 1-month-old. (B) 2-month-old. Circled areas: SNc. Scale bars = 20 µm.

